# piRNAs prevent runaway amplification of siRNAs from ribosomal RNAs and histone mRNAs

**DOI:** 10.1101/2020.06.15.153023

**Authors:** Brooke E. Montgomery, Tarah Vijayasarathy, Taylor N. Marks, Kailee J. Reed, Taiowa A. Montgomery

**Author notes:** Authors contributed equally (Brooke E. Montgomery, Tarah Vijayasarathy).

## Abstract

Piwi-interacting RNAs (piRNAs) are a largely germline-specific class of small RNAs found in animals. Although piRNAs are best known for silencing transposons, they regulate many different biological processes. Here we identify a role for piRNAs in preventing runaway amplification of small interfering RNAs (siRNAs) from certain genes, including ribosomal RNAs (rRNAs) and histone mRNAs. In *Caenorhabditis elegans*, rRNAs and some histone mRNAs are heavily targeted by piRNAs, which facilitates their entry into an endogenous RNA interference (RNAi) pathway involving a class of siRNAs called 22G-RNAs. Under normal conditions, rRNAs and histone mRNAs produce relatively low levels of 22G-RNAs. But if piRNAs are lost, 22G-RNA production is highly elevated. We show that 22G-RNAs produced downstream of piRNAs likely function in a feed-forward amplification circuit. Thus, our results suggest that piRNAs facilitate low-level 22G-RNA production while simultaneously obstructing the 22G-RNA machinery to prevent runaway amplification from certain RNAs. Histone mRNAs and rRNAs are unique from other cellular RNAs in lacking polyA tails, which may promote feed-forward amplification of 22G-RNAs. In support of this, we show that the subset of histone mRNAs that contain polyA tails are largely resistant to silencing in piRNA mutants.

## INTRODUCTION

Piwi-interacting RNAs (piRNAs) affect many different processes in the germline (1–3). Perturbing the piRNA pathway often leads to sterility, in part because of its critical role in silencing transposable elements (1). In the nematode *Caenorhabditis elegans*, loss of piRNAs does not lead to immediate sterility, but instead causes a gradual loss of fertility over numerous generations, such that the germline loses its immortal nature (4). Whether germline mortality in *C. elegans* piRNA mutants is due to activation of transposons and other repetitive elements has not been resolved. Several alternative models have been proposed that could also explain the progressive sterility of piRNA mutants. For example, we and others showed that piRNAs prevent silencing of essential genes by an endogenous RNAi pathway involving small interfering RNAs (siRNAs) (5, 6). Recently, the germline mortality of piRNA mutants was linked specifically to the silencing of essential histone genes (7).

In *C. elegans*, piRNAs associate with a single Piwi protein, PRG-1, where they act as sequence-specific guides to direct the complex to target mRNAs (8–10). The interaction between piRNAs and target mRNAs in some instances leads to the production and subsequent amplification of siRNAs called 22G-RNAs by an RNA-dependent RNA polymerase in association with a collection of proteins called the Mutator complex (11–14). The Mutator complex is seeded by MUT-16 adjacent to the germ granules that house the piRNA machinery (8, 13). This is the same 22G-RNA pathway that is activated by canonical siRNAs produced from double-stranded RNA during exogenous RNA interference (RNAi) and is therefore commonly referred to as endogenous RNAi (11). piRNA-dependent 22G-RNAs bind a worm-specific class of Argonautes called WAGOs. A genetically distinct class of 22G-RNAs bind to the Argonaute CSR-1 and is thought to act in opposition to piRNAs to promote or fine-tune gene expression (15–21).

*C. elegans* contains thousands of distinct piRNAs that require only partial sequence complementarity to bind target mRNAs (22–24). Consequently, piRNAs engage most if not all mRNAs in the germline (22, 23, 25). Furthermore, the 22G-RNAs produced from piRNA targets are thought to serve as a memory of piRNA activity that can persist in the absence of the initial piRNA trigger (26–28). Because of these features, identifying the precise mechanism by which piRNAs regulate gene expression is challenging. For example, it is unclear if piRNAs physically direct mRNAs into the RNAi pathway or if it is an indirect consequence of an RNA being retained in germ granules.

While loss of piRNAs leads to a reduction in WAGO-class 22G-RNA levels from many mRNAs in the germline, core histones instead produce elevated levels (7, 29). In this study, we find that ribosomal RNAs also hyperaccumulate 22G-RNAs in *prg-1* mutants. 22G-RNA amplification from both rRNAs and histone mRNAs is dependent on the WAGO-class 22G-RNAs produced downstream of piRNAs. Thus, piRNAs are required both for the production of WAGO-class 22G-RNAs and to prevent their overamplification from certain RNAs. Our results suggest that piRNAs facilitate binding of the RNAi machinery and subsequent 22G-RNA production, while also suppressing 22G-RNA feed-forward amplification, possibly by obstructing access to the RNAi machinery. Histone mRNAs and rRNAs are distinct from other cellular RNAs in that they lack polyA tails (30). We propose that polyadenylation helps to protect RNAs from runaway 22G-RNA amplification.

## RESULTS

### Hyperproduction of 22G-RNAs from histones and other coding genes in *prg-1* mutants

We and others recently showed that histone transcripts are directed into an endogenous RNAi pathway in *piwi/prg-1* mutants where they spawn high levels of 22G-RNAs, which is in some instances coincident with reduced histone mRNA levels (7, 29). To determine if this phenomenon is unique to histones, we examined 22G-RNA production from all annotated protein coding genes using RNA-seq datasets from dissected distal gonads from wild type and *prg-1* and *mut-16* mutants (29). The majority of protein coding genes yielded 22G-RNA reads in wild type animals, and to a lesser extent in *prg-1* and *mut-16* mutants (Fig. 1A). A small subset of genes produced highly elevated levels of 22G-RNAs in *prg-1* mutants, but not in *mut-16* mutants (Fig. 1B-1C). We identified 222 genes that yielded >250 normalized reads (~30 reads per million total, RPM) in *prg-1* mutants and which were upregulated >5 fold in *prg-1* mutants relative to wild type animals. These arbitrarily applied cutoffs captured most histones and minimized what may be noise in 22G-RNA expression (Fig. 1B). Of the 222 genes, ~30% were detectably downregulated >1.3 fold at the mRNA level (Fig. 1D). The majority of the 66 genes (i.e. 39) that produced elevated levels of 22G-RNAs and reduced mRNA levels are not histones, for example, the aldehyde dehydrogenase *alh-7* (Fig. 1D-1E).

**Figure 1.**
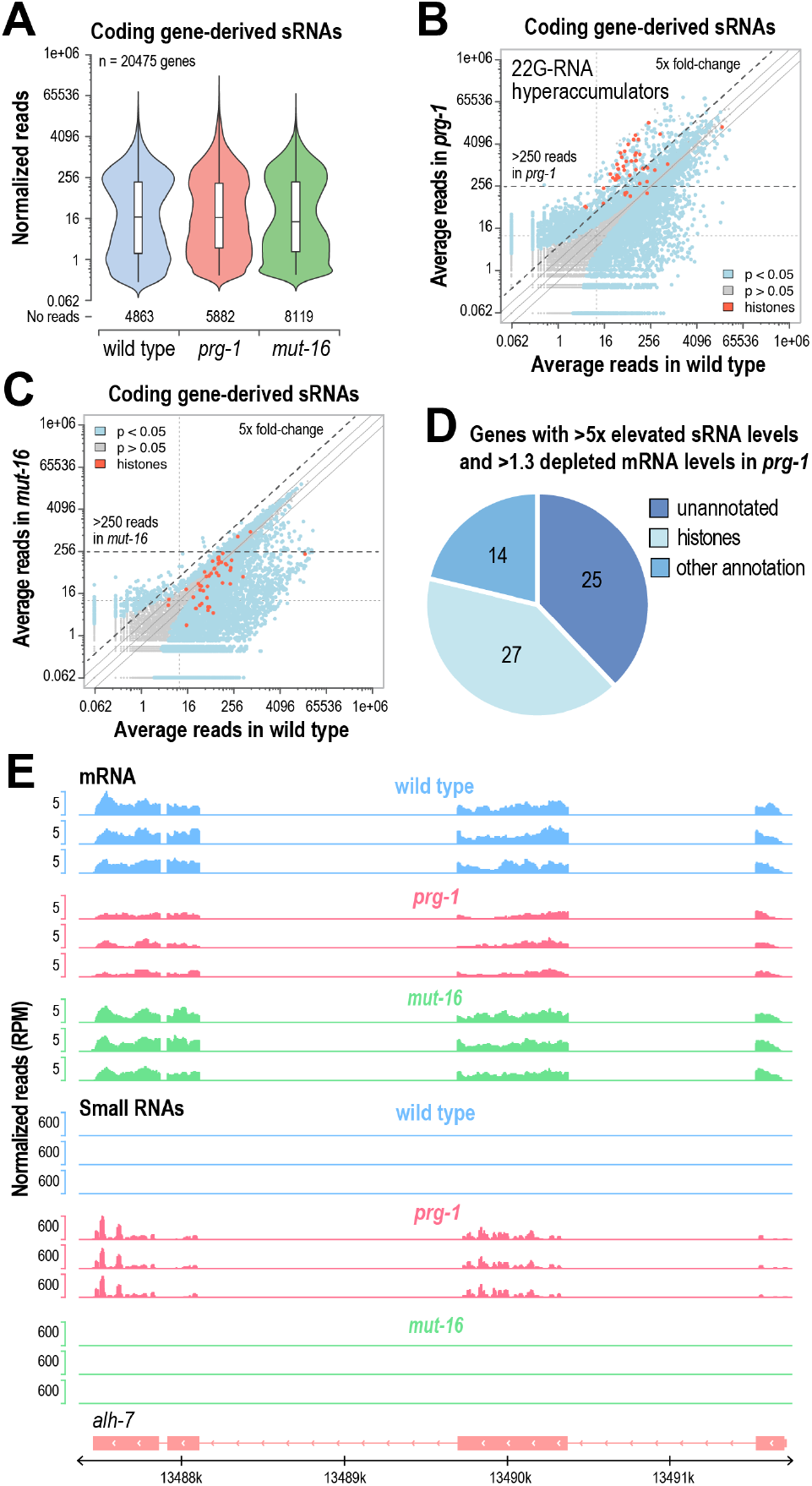
Hyperaccumulation of 22G-RNAs in *prg-1* mutants. (A) Normalized small RNAs reads from annotated coding genes in small RNA-seq libraries from dissected gonads of wild type, *prg-1(n4357),* and *mut-16(pk710)* mutants (n=3 biological replicates for each strain). (B-C) Scatter plots displaying each gene as a function of normalized small reads in wild type and *prg-1(n4357)* (B) or *mut-16(pk710)* (C) mutants. Libraries from dissected distal gonads (n=3 biological replicates for each strain). (D) Pie chart classifying genes that produced >5-fold as many 22G-RNA reads in *prg-1(n4357)* mutants compared to wild type and for which the corresponding mRNA was downregulated >1.3-fold. (E) mRNA and small RNA read distribution across a representative gene locus, *alh-7*. 3 biological replicates are shown for wild type, *prg-1(n4357*), and *mut-16(pk710)*.

### 22G-RNA amplification from ribosomal RNAs in *prg-1* mutants

We next examined small RNA production from long non-coding RNAs and structural RNAs, including snRNAs, scRNAs, snoRNAs, tRNAs, and rRNAs, in *prg-1* mutants to identify if such features are also targeted for hyperproduction of 22G-RNAs in the absence of piRNAs. Many non-coding RNAs produced relatively high levels of 22G-RNAs, a subset of which were depleted in *prg-1*, as well as in *mut-16*, indicating that they are targeted by piRNAs and routed into the RNAi pathway, similar to many coding genes (Fig. 2A-2B). However, rRNAs stood out as having high levels of small RNAs that were elevated >5 fold in *prg-1* mutants, but were not substantially affected in *mut-16* mutants (Fig. 2A-2B). These small RNAs bare the hallmarks of 22G-RNAs: antisense orientation, preference for a 5’G, and length of 22-nts, and are thus likely the ribosomal siRNAs (risiRNAs) previously associated with rRNA misprocessing, although we have no reason to believe that rRNAs are misprocessed in *prg-1* mutants (Fig. 2C-2D) (31, 32).

**Figure 2.**
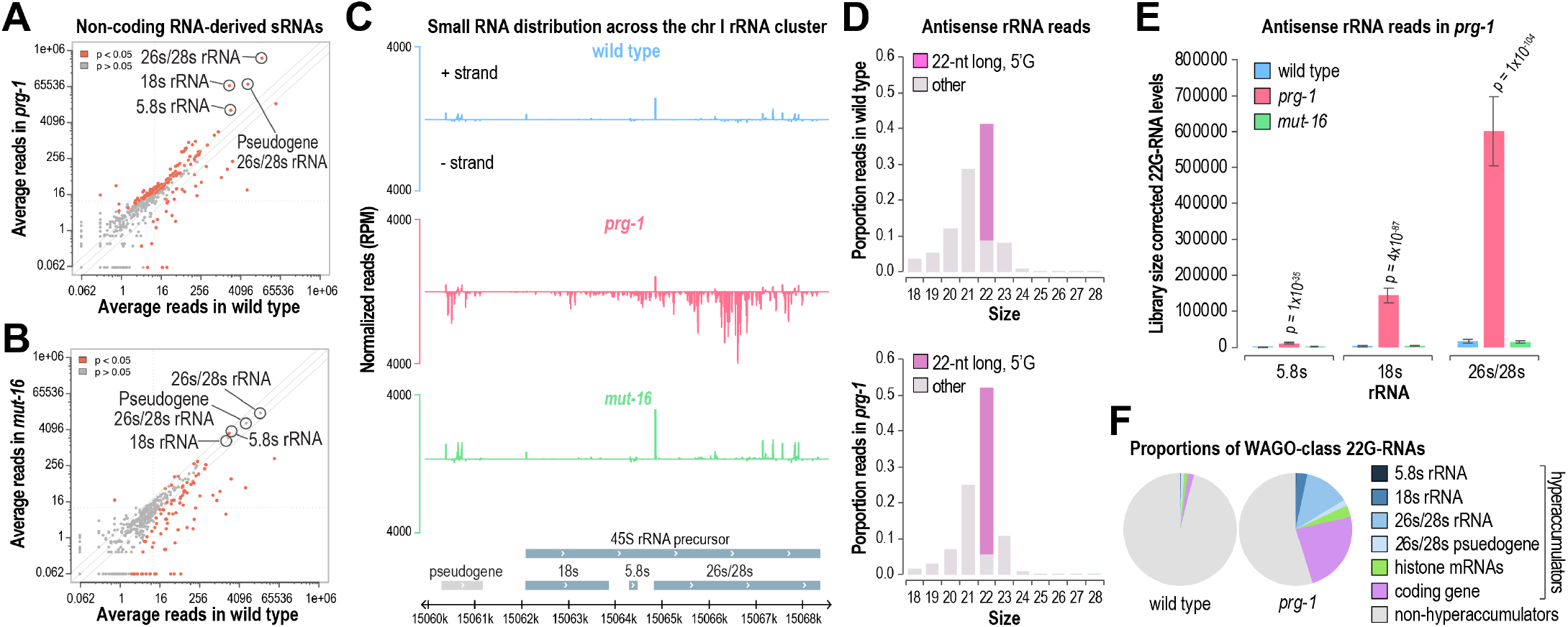
Ribosomal RNAs hyperaccumulate 22G-RNAs in *prg-1* mutants. (A-B) Scatter plots displaying each non-coding RNA as a function of normalized small reads in wild type and *prg-1(n4357)* (A) or *mut-16(pk710)* (B) mutants. rRNAs are circled. Libraries from dissected distal gonads (n=3 biological replicates for each strain). (C) Small RNA read distribution across an rRNA locus. 1 of 3 biological replicates is shown for wild type, *prg-1(n4357*), and *mut-16(pk710)*. (D) Antisense rRNA-derived small RNA read size distribution in wild type and *prg-1(n4357*) mutants. 22G-RNAs are colored purple. (E) Normalized rRNA-derived small RNA reads in wild type, *prg-1(n4357*), and *mut-16(pk710)*. Standard deviation for 3 biological replicates is shown for each strain. (F) Pie chart displaying the proportion of all WAGO-class 22G-RNA reads from features classified as 22G-RNA hyperaccumulators (>5-fold 22G-RNA levels in *prg-1* mutants relative to wild type) or non-hyperaccumulators. WAGO-class features were expanded to include the hyperaccumulators that are not classified as WAGO targets.

The 22-nt, 5’G bias among rRNA-derived small RNAs was more prominent in *prg-1* mutants than in wild type. This may reflect a shift to production via the endogenous RNAi pathway in *prg-1* mutants, as we previously found that WAGO-class 22G-RNAs have a stronger preference for 22-nt species than CSR-1-class 22G-RNAs, which are produced through an alternative pathway (Fig. 2D) (16, 33). As noted above, rRNA-derived 22G-RNAs are not depleted in *mut-16* mutants, suggesting that the majority are indeed produced through the CSR-1 pathway under normal conditions (Fig. 2B). We propose that rRNAs are routed primarily into the CSR-1 pathway for low-level 22G-RNA production and gene licensing and that in *prg-1* mutants, the balance shifts to WAGO-class 22G-RNA production. Consistent with this possibility, rRNA-derived 22G-RNAs, were previously shown to bind both CSR-1 and WAGO-1 (32).

5.8s rRNA-derived 22G-RNAs levels were elevated ~5-fold and 18s- and 26s/28s rRNA-derived 22G-RNAs levels were elevated >30-fold in *prg-1* mutants (Fig. 2E). 22G-RNAs produced from rRNAs, histones, and other annotated coding genes that displayed hyperaccumulation in *prg-1* mutants accounted for ~45% of all WAGO-class 22G-RNA reads in *prg-1* mutants, but only a very small fraction in wild type animals (Fig. 2F). This points to substantial repurposing of the 22G-RNA machinery in the absence of piRNAs. Disabling a single class of primary small RNAs that trigger WAGO-class 22G-RNA production (i.e. ERGO-1-class 26G-RNAs) enhances exogenous RNAi, likely because of competition for shared factors, suggesting that RNAi functions at or near capacity (34). Therefore, the dramatic shift in WAGO-class 22G-RNA production to rRNAs, histones, and other coding genes may indirectly impact 22G-RNA production across a large number of WAGO targets. Alternatively, hyperproduction of 22G-RNAs may be facilitated by loss of other 22G-RNAs in *prg-1* mutants.

### Hyperproduction of 22G-RNAs due to runaway amplification in *prg-1* mutants

Hyperaccumulation of 22G-RNAs in *prg-1* mutants could be related to a direct role for piRNAs in suppressing 22G-RNA production. Alternatively, it could be due to loss of piRNA-dependent 22G-RNAs, which may function within a self-reinforcing feedback loop (12, 14). To distinguish between these possibilities, we analyzed small RNA high-throughput sequencing data from three strains (6): 1) A control strain generated by crossing two strains carrying distinct genetic causes for loss of WAGO-class 22G-RNAs and then selecting for animals homozygous wild type for each gene that was mutant in the parent strains (Fig. 3A, top). In this strain, the WAGO-class 22G-RNA pathway is reanimated from ancestral lines that lacked the ability to produce WAGO-class 22G-RNAs. This strain has both piRNAs and piRNA-dependent 22G-RNAs and is effectively wild type. 2) A strain in which the WAGO-class 22G-RNA pathway was reestablished as in 1 but with the male parent containing a mutation in *prg-1*. F2 animals homozygous for the *prg-1* mutation but otherwise wild type were selected (Fig. 3A, middle). This strain retains the maternal heritable 22G-RNAs produced downstream of *prg-1* that are thought to provide a memory of piRNA activity, but lacks piRNAs. This strain is effectively *prg-1−/−*, but otherwise wild type. 3) A strain generated as above but with both parents containing a mutation in *prg-1*. Progeny wild type for each RNAi factor but homozygous mutant for *prg-1* were selected. In this strain, both piRNAs and piRNA-dependent 22G-RNAs are lost but WAGO-class 22G-RNA production is otherwise restored to normal (Fig. 3A, bottom). Note that this strain was generated on *mut-16* RNAi to suppress reactivation of the WAGO-class 22G-RNA pathway, which would otherwise lead to immediate sterility in the absence of piRNAs (6). After homozygosing the 3 strains, they were subjected to RNA isolation and small RNA-sequencing.

**Figure 3.**
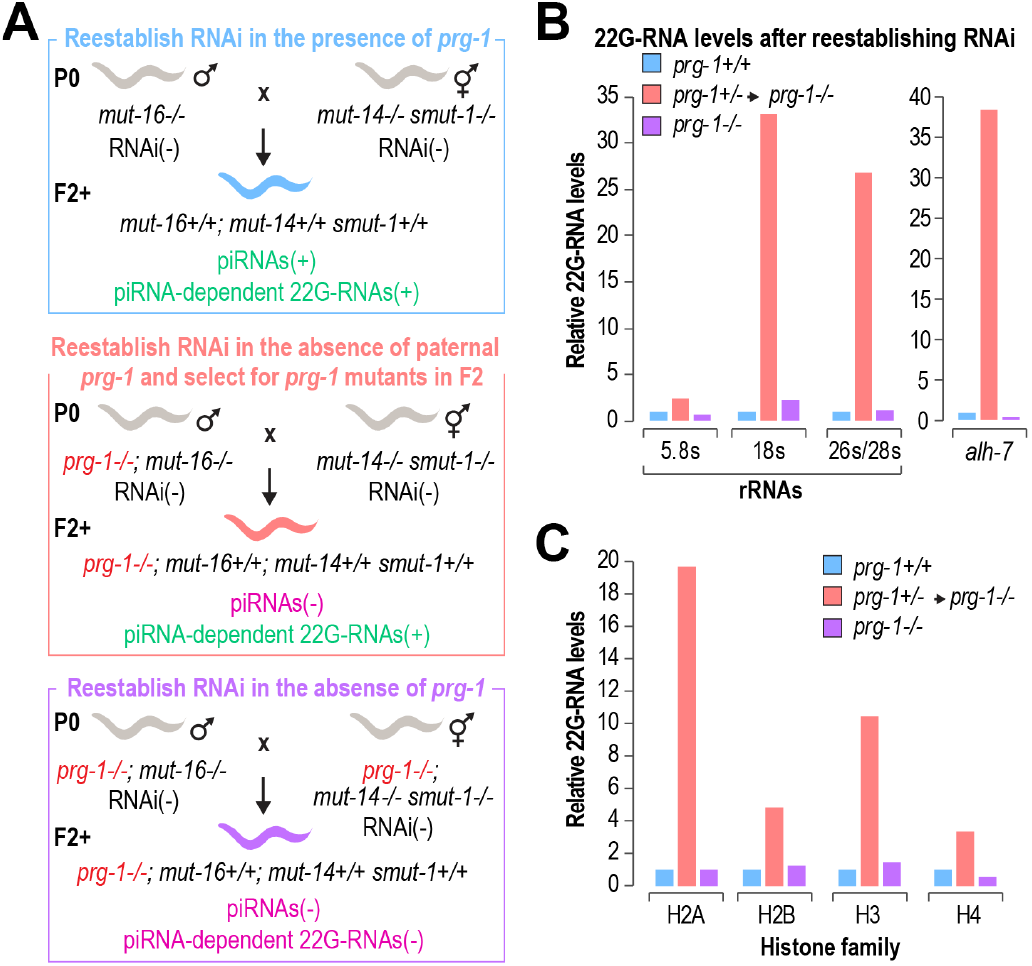
*prg-1* prevents runaway amplification of 22G-RNAs. (A) A genetics-based approach for reestablishing RNAi in the presence or absence of *prg-1*. RNAi/22G-RNA competency in the parent strains is indicated by RNAi(+) or RNAi(-). Presence or absence of piRNAs and piRNA-dependent 22G-RNAs in the progeny of the crosses is similarly indicated. Alleles: *prg-1(n4357), mut-16(pk710)*, and *mut-14(mg464) smut-1(tm3101)*. (B-C) Relative levels of 22G-RNAs derived from rRNAs and *alh-7* (B) and histones (C) after reestablishing 22G-RNA production in the presence or absence of *prg-1* (no biological replication). The bars in B and C are colored-matched to the worms in the diagrams in A.

Disregarding the complex genetics used to generate the strains above, they have following attributes: 1) piRNA(+)/piRNA-dependent 22G-RNA(+), 2) piRNA(-)/piRNA-dependent 22G-RNA(+), and 3) piRNA(-)/piRNA-dependent 22G-RNA(-). Using the three strains we could disentangle the roles of piRNAs from that of piRNA-dependent 22G-RNAs in the hyperaccumulation of 22G-RNAs. In the piRNA(-)/piRNA-dependent 22G-RNA(+) strain, 22G-RNA production from rRNAs and histones was strongly upregulated relative to the *prg-1+/+* (wild type) control (Fig. 3B-3C, pink vs blue worms and bars). This was to be expected given that the strain is more or less identical to the *prg-1−/−* strains used in the original experiments. In contrast, in the piRNA(-)/piRNA-dependent 22G-RNA(-) strain, 22G-RNA production from rRNAs and histones was not upregulated relative to the *prg-1*+/+ (wild type) control (Fig. 3B-3C, purple vs blue worms and bars). Similarly, 22G-RNA production from *alh-7* was also specifically upregulated in the piRNA(-)/piRNA-dependent 22G-RNA(+) strain (Fig. 3B). Thus, piRNA-dependent 22G-RNAs are required for the hyperproduction of 22G-RNAs in *prg-1* mutants. This points to a model in which PRG-1-piRNA complexes facilitate low-level WAGO-class 22G-RNA production from rRNAs, histones, and other coding genes, while at the same time preventing feed-forward amplification.

### Polyadenylated histone mRNAs evade runaway amplification in *prg-1* mutants

We were not able to identify a single common feature that might explain why some RNAs are prone to runaway amplification of 22G-RNAs in *prg-1* mutants. However, histones and rRNAs, which account for roughly half of the 22G-RNAs that hyperaccumulate in *prg-1* mutants (Fig. 2F), are unique from other RNAs in that they lack polyA tails (30). The Argonaute CSR-1 was previously implicated in histone mRNA 3’-end processing and it was recently shown that in the absence of *prg-1*, CSR-1 associates more strongly with WAGO-1, which may promote 22G-RNA amplification (7, 35). More generally, however, it is possible that non-polyadenylated RNAs are hypersusceptible to runaway amplification, which may or may not be facilitated by CSR-1. To explore this possibility, we first identified which mRNAs are polyadenylated in *C. elegans* by subjecting rRNA-depleted total RNA to polyA selection or a control treatment and then performed high-throughput mRNA sequencing (Fig. 4A). The majority of histone mRNAs were dramatically enriched in the non-polyA-selected mRNA libraries relative to the polyA-selected libraries, as predicted because of their lack of polyA tails (Fig. 4B) (36). The linker histone H1 and the variant H3.3 were not enriched in the non-polyA-selected libraries, consistent with previous studies demonstrating that, unlike the canonical core histones, they do possess polyA tails (Fig. 4B) (37). Within each of the core histone families (H2A, H2B, H3, and H4), a single member was not enriched in the non-polyA-selected library relative to the polyA-selected library (Fig. 4B). However, the H3 representative was only modestly enriched compared to other histones and is predicted to produce a truncated protein, whereas H2B had two members, one of which was weakly expressed (Fig. 4B).

**Figure 4.**
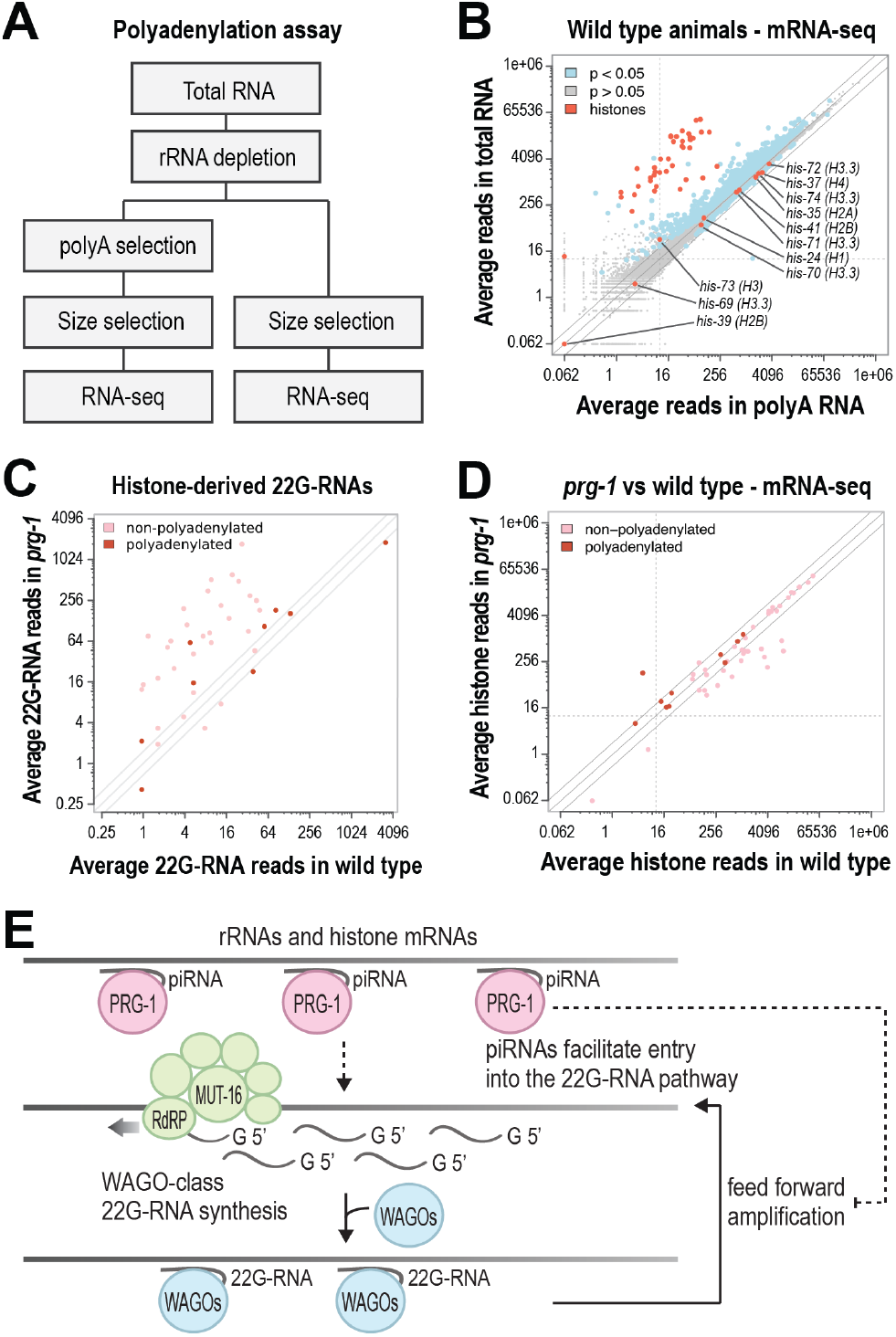
Polyadenylated histone mRNAs are less susceptible to silencing in *prg-1* mutants. (A) Schematic depicting a high-throughput sequencing-based polyadenylation assay. (B) Scatter plot displaying each gene as a function of normalized reads from total rRNA-depleted RNA and rRNA-depleted, polyA-selected RNA. Note that the original RNA was the same for each but was treated as diagramed in A. (Libraries from whole adult animals, n=3 biological replicates). (C) Scatter plot displaying each histone as a function of normalized 22G-RNA reads in wild type and *prg-1(n4357)* mutant gonads (n=3 biological replicates). Only uniquely mapping reads are included. (D) Scatter plot displaying each histone as a function of normalized mRNA reads in wild type and *prg-1(n4357)* mutant gonads (n=3 biological replicates). (E) Model for 22G-RNA runaway amplification. PRG-1-piRNA complexes bind to rRNAs and histones which facilitates entry into the WAGO-class 22G-RNA pathway. PRG-1 then suppresses feed-forward amplification of 22G-RNAs.

Having determined that some histone mRNAs likely contain polyA tails, we assessed whether this small subset of histone genes was downregulated in *prg-1* mutants similarly to non-polyadenylated histone mRNAs. Each of the mRNAs enriched in non-polyA mRNA libraries was unchanged or only slightly misregulated in *prg-1* mutants (Fig. 4C). Additionally, the polyadenylated histones were less prone to hyperproduction of 22G-RNAs in *prg-1* mutants (Fig. 4C). However, many histone mRNAs that were upregulated in our non-polyA-selected libraries displayed only modest upregulation of 22G-RNAs and were not substantially downregulated in *prg-1* mutants (Fig. 4C-4D). Because of sequence similarity between histone genes within the same family, it is likely that 22G-RNAs produced from one histone gene target multiple paralogous histones, which may confound our analysis to some extent. Nonetheless, these results are consistent with the lack of a polyA tail promoting runaway 22G-RNA amplification.

The polyadenylated histones also contain introns, which are absent in the histones, as well as in rRNAs, that hyperaccumulate 22G-RNAs. However, most of the other genes that we classified as 22G-RNA hyperaccumulators do contain introns, suggesting that introns do not protect against runaway amplification in *prg-1* mutants. Nonetheless, there are certainly other features that contribute to runaway amplification, as most non-histone genes that were highly upregulated in our non-polyA selected versus polyA selected libraries were not identified as hyperaccumulators (Fig. 4B).

## DISCUSSION

Our results point to a model in which piRNAs bind to RNA targets to facilitate the production of 22G-RNAs, while at the same time preventing overamplification via a self-enforcing feedback loop (Fig. 4E). This phenomenon was primarily associated with histones and rRNAs, but it may occur much more broadly, as it is difficult to tease out the contribution of piRNAs in facilitating 22G-RNA production from their role in preventing 22G-RNA overamplification. The absence of polyA tails on rRNAs and histone mRNAs may contribute to runaway amplification of 22G-RNAs. For example, polyA-binding proteins may limit access to the 22G-RNA machinery or direct them out of germ granules for translation. Alternatively, mRNAs that lack polyA tails may be primed to serve as templates for 22G-RNA production. Within the Mutator complex that produces 22G-RNAs is a protein, RDE-3, that adds polyU tails to target mRNAs, but the mRNAs are first stripped of their polyA tails by either cleavage or 3’-5’ trimming (38). This processing step may not be required for RNAs that already lack polyA tails and could therefore accelerate 22G-RNA production.

rRNAs and histone mRNAs are heavily targeted by PRG-1-piRNA complexes across much of their length (23). PRG-1-piRNA complexes may physically impede association with the 22G-RNA machinery to prevent runaway amplification under normal conditions. This hints at an alternative function for piRNAs: rather than physically directing mRNAs into the 22G-RNA pathway, piRNAs may act as a molecular sieve of sorts that temporarily traps RNAs in germ granules. Presumably the trapped RNAs are acted on by gene-licensing factors, such as CSR-1, or gene-silencing factors, such as the WAGO Argonautes, or both. This would help to explain why essentially all germline mRNAs are targeted by piRNAs, while only a subset are silenced (23, 29, 39). The sheer abundance of rRNAs and histone mRNAs may overwhelm the cellular machinery that processes them. Without PRG-1 and polyA-binding proteins to help protect rRNAs and histones, 22G-RNAs may engage in a feedback loop that leads to runaway amplification by RNA-dependent RNA polymerases. This model may explain why hundreds of CSR-1 targets are also depleted of 22G-RNAs in *prg-1* mutants (29). Perhaps a similar molecular sieve model could help to explain the highly abundant pachytene piRNAs found in mice, which also have expansive targeting capacity but lack a clear role in regulating gene expression (40).

## METHODS

### Strains

The SX922[prg-1(n4357) I], NL1810[mut-16(pk710) I], and GR1948[mut-14 (mg464) smut-1 (tm3101) V] strains were described by Das et al. (9), Vastenhouw et al. (41), and Philips et al. (33) respectively.

### RNA isolation

RNA isolation from gonads was done in Reed et al. (29). Worms used for RNA isolation in this study were washed three times in M9 buffer, flash frozen in liquid nitrogen, and lysed in Trizol (LifeTechnologies). RNA was isolated using two rounds of chloroform extraction followed by isopropanol precipitation or using the Direct-zol RNA Miniprep Plus kit (Zymo Research).

### mRNA-seq libraries

mRNA-seq libraries were prepared using the NEBNext Ultra II Directional RNA Library Prep Kit for Illumina (NEB) as described (29), or the TruSeq Stranded Total RNA Library Prep Human/Mouse/Rat kit (Illumina) following the manufacturer’s protocol. Prior to library preparation, rRNA was depleted using the Ribo-Zero rRNA Removal Kit (Illumina). Where indicated in the results, polyadenylated RNAs were selected using the NEBNext Poly(A) mRNA Magnetic Isolation Module (NEB) after rRNA depletion. Samples were sequenced on an Illumina NextSeq 500 (High Output Kit, single-end, 75 cycles).

### mRNA-seq data analysis

Data analysis was done as described (29). Briefly, quality filtering was done with Trimmomatic v. 0.35 (42). Reads were mapped to the *C. elegans* genome (Wormbase release WS230) using Star v. 2.5.0a (43). Reads for each feature were counted using RSEM v. 1.3.0 (44). Differential expression analysis was done using DESeq2 v. 1.18.1 (45). Plots were drawn in R, Excel, and IGV (46).

### sRNA-seq libraries

Small RNA libraries were prepared as described (29). Briefly, 16-30-nt RNAs were size selected on 17% denaturing polyacrylamide gels and treated with RNA polyphosphatase (Illumina) to reduce 5′ di- and triphosphates to monophosphates to enable 5′ adapter ligation. Sequencing libraries were prepared with the NEBNext Multiplex Small RNA Library Prep Set for Illumina (NEB). Libraries were size selected on 10% polyacrylamide gels and sequenced on an Illumina NextSeq 500 (High Output Kit, single-end, 75 cycles).

### sRNA-seq data analysis

Small RNA data analysis was done as described (29). Briefly, small RNAs were parsed from adapters, quality filtered, and mapped to the *C. elegans* genome (Wormbase release WS230) using CASHX v. 2.3 (47). Imperfectly matching reads were not included in our analysis. Reads from specific features were counted using custom Perl scripts (48). Differentially expressed small RNAs were identified using DESeq2 v. 1.18.1 (45). R, Excel, and IGV (46) were used for drawing plots.

### Reestablishing WAGO-class 22G-RNA production

WAGO-class 22G-RNA production was reestablished in RNAi-defective mutants using the genetics approach in Fig. 3A, as described (6). Because reestablishing WAGO-class 22G-RNA production in the absence of *prg-1* causes sterility, *mut-16* was inactivated by RNAi during the cross in which both parents were *prg-1* mutant while homozygosing and expanding the strain. Afterwards, the animals were transferred off of *mut-16* RNAi to reestablish the RNAi pathway prior to RNA isolation for small RNA sequencing (6). The other strains were grown in parallel but were not treated with *mut-16* RNAi.

## ACKNOWLEDGEMENTS

High-throughput sequencing was done at the Colorado State University Next Generation Sequencing Facility by Mark Stenglein and Marylee Layton. Thanks to Carolyn Phillips, René Ketting, and Germano Cecere for insightful comments and suggestions. The NL4415 [henn-1(pk2295)] strain was a kind gift from René Ketting. Additional strains were provided by the CGC, which is funded by the NIH (P40 OD010440). This work was supported by the NIH [R35GM119775 to T.A.M. and T32GM132057 to K.J.R.].

## Notes

### Competing Interest Statement

The authors have declared no competing interest.

